# Deep tissue scattering compensation with three-photon F-SHARP

**DOI:** 10.1101/2021.08.18.456049

**Authors:** Caroline Berlage, Malinda Tantirigama, Mathias Babot, Diego Di Battista, Clarissa Whitmire, Ioannis N. Papadopoulos, James F. A. Poulet, Matthew Larkum, Benjamin Judkewitz

## Abstract

Optical imaging techniques are widely used in biological research, but their penetration depth is limited by tissue scattering. Wavefront shaping techniques are able to overcome this problem in principle, but are often slow and their performance depends on the sample. This greatly reduces their practicability for biological applications. Here we present a scattering compensation technique based on three-photon (3P) excitation, which converges faster than comparable two-photon (2P) techniques and works reliably even on densely labeled samples, where 2P approaches fail. To demonstrate its usability and advantages for biomedical imaging we apply it to the imaging of dendritic spines on GFP-labeled layer 5 neurons in an anesthetized mouse.

## Introduction

Fluorescence-based microscopy methods have become a standard approach to visualize biological processes from the scale of individual molecules up to entire organisms. Nonetheless, tissue turbidity limits their *in vivo* application to superficial areas or small animals (1).

As light travels through inhomogeneous media such as biological tissue, it deviates from its straight path due to refractive index variations. The scattering of the excitation light degrades microscopy images in two ways: First, scattered photons excite out-of-focus fluorescence, thus increasing the background. Second, scattering leads to lower light intensity at the desired focal spot and therefore reduces the signal. Together, these two effects decrease the signal-to-background ratio (SBR), making it harder to resolve the structures of interest at increasing imaging depths.

Several methods have been developed to address the first problem of out-of-focus excitation, thus reducing the contribution of scattered light. In confocal microscopy (2), a pinhole on the detection side rejects light emitted away from the focus. Alternatively, nonlinear techniques such as multiphoton microscopy (3) require the coincident arrival of more than one photon to excite a fluorophore and thus limit the excitation to the vicinity of the focal point. However, both techniques rely on ballistic (non-scattered) light. Because the amount of ballistic photons decreases exponentially with propagation distance, the imaging depth for these methods remains limited (1, 4).

An alternative approach addresses both issues simultaneously by shaping the incident wavefront to compensate for the sample-induced aberrations, which can effectively refocus light to a diffraction-limited point. This increases the signal and reduces background at the same time. By introducing a compensatory distortion of the incoming light field before it enters the tissue, the net aberration can be minimized and a diffraction-limited focus can be formed inside the tissue.

Inspired by astronomy (5), adaptive optics (AO) microscopy uses wavefront shaping devices such as deformable mirrors or spatial light modulators (SLMs) to modulate the light phase at the back focal plane of the objective to compensate for aberrations introduced by the sample. However, the main challenge is to determine the aberrations introduced by the sample, so the correction pattern can be determined. In living organisms, this is further complicated by the lack of direct access to the focal plane or transmitted light.

Multiple AO techniques have been developed towards this aim, which can be divided into two broad categories (6, 7): The correction pattern can either be determined directly, often using a Shack-Hartmann wavefront sensor (“direct AO”) (8, 9), or without an additional sensor (“indirect AO”), for example by iteratively optimizing image metrics such as brightness (10–12) or by sequentially measuring aberrations for different pupil segments (13, 14).

While indirect AO methods are often easier to implement because they do not rely on an additional sensor, they are often slow, taking seconds to minutes to determine the correction pattern. This limits their application *in vivo*. Direct AO methods are often faster, but only work in weakly scattering samples and require a bright guide star to measure the wavefront, which often exceeds the photon budget in living samples (7). These AO methods are mainly used to correct for low-order aberrations, for example due to sample surface curvature or slowly varying tissue properties. However, many regions of biological interest are deep inside tissue, so that light also accumulates high-order aberrations due to scattering.

Fortunately, even in highly scattering regimes inside biological tissue, wavefront shaping can achieve a diffraction-limited focus (15–17), as the effects of absorption in biological tissue are negligible (18) and most of the light is scattered in the forward direction (19–22). However, because of the highly complex distributions of scatterers within a biological medium, necessary correction patterns contain many modes and require a large amount of computational or experimental time to determine. At the same time, the spatial distribution of scatterers is dynamic and can decorrelate at short time scales (23). Hence, the main challenge in deep tissue imaging is to find a way to quickly determine correction patterns.

Different guidestar-based methods have been developed (24), which use fluorescent targets such as beads (25), photoacoustic feedback (26) or ultrasound (27–29) to determine the wavefront correction map. An alternative approach is to determine the correction by modulating the SLM pixels to iteratively improve the correction pattern (30–33). However, the measurement speed of this approach is limited by the modulation speed of the SLM. Both of these approaches suffer from slow speed or require a transmission geometry that makes their *in vivo* use challenging.

To decouple the measurement from the SLM speed, focus scanning holographic aberration probing (F-SHARP) (34, 35) uses non-linear excitation by two interfering beams to directly measure both amplitude and phase of the scattered light distribution inside the tissue. This knowledge can then be used to non-invasively correct for the measured deviations from a diffraction-limited PSF.

In this work, we extend F-SHARP to three-photon (3P) excitation. We show that F-SHARP not only benefits from the increased penetration depth and signal-to-background ratio of 3P excitation (36), but that the third-order non-linearity leads to qualitatively different performance. First, we show that unlike two-photon (2P) techniques, 3P F-SHARP does not rely on sample contrast to converge, and works reliably even on samples with homogeneous three-dimensional distribution of fluorescence. Additionally, the iterative optimization of 3P F-SHARP converges faster than that of 2P F-SHARP. This means that it requires fewer iterations, and thus less measurement time. We then demonstrate the applicability to *in vivo* microscopy by imaging spines of layer 5 pyramidal neurons in mice.

## Principle of operation

In laser-scanning microscopy, an image is acquired by scanning a focused beam across the sample. The spatial profile of this beam at the focal plane is defined as the intensity pointspread function *I*_PSF_. For linear fluorescence microscopy, the acquired image then consists of a convolution of the fluorescent sample features with *I*_PSF_.

While conventional detectors only measure light intensity, the beam propagation depends on both amplitude and phase of the complex electric field. We call the complex equivalent of *I*_PSF_ the electric field point spread function *E*_PSF_ or E-field PSF, with *I*_PSF_ = |*E*_PSF_ |^2^. In an ideal case (assuming a circular aperture), the *I*_PSF_ is a diffraction-limited Airy function, with most of its energy located in the central lobe. However, as described in the introduction, light propagation through a disordered medium such as biological tissue degrades the PSF via scattering and aberrations. In the following, we will use the term ’scattered *E*_PSF_’ to describe any E-field PSF that deviates from an ideal one, including both aberrations and scattering.

Multiphoton microscopy takes advantage of nonlinear excitation. Mathematically, this image formation can be described as a convolution operation of the object with higher powers of the *I*_PSF_ (e.g. |*I*_PSF_|^3^ for 3P microscopy). This reduces fluorescence excitation due to the weaker side lobes of the scattered E-field PSF. However, the power contained in the central peak of the PSF still remains reduced.

The aim of F-SHARP is to not only suppress out-of-focus excitation, but to refocus the full power back into a diffraction-limited spot. Towards this aim, F-SHARP directly measures the complex scattered *E*_PSF_ inside the scattering medium. Using this reconstructed E-field, deviations from the ideal, diffraction-limited shape can be corrected by applying a correction pattern onto a spatial light modulator (SLM). With this corrected wavefront, the sample can then be imaged with increased signal-to-noise ratio (SNR).

F-SHARP estimates the scattered *E*_PSF_ without direct access to the focal plane, and with detectors only capable of measuring intensity and not phase. To do so, the excitation light is divided into two beams. A “strong beam” which contains most of the excitation power is parked in the center of the field of view (FOV) and a “weak beam” is scanned across the strong beam to probe its shape (Fig. 1b).

**Fig. 1.**
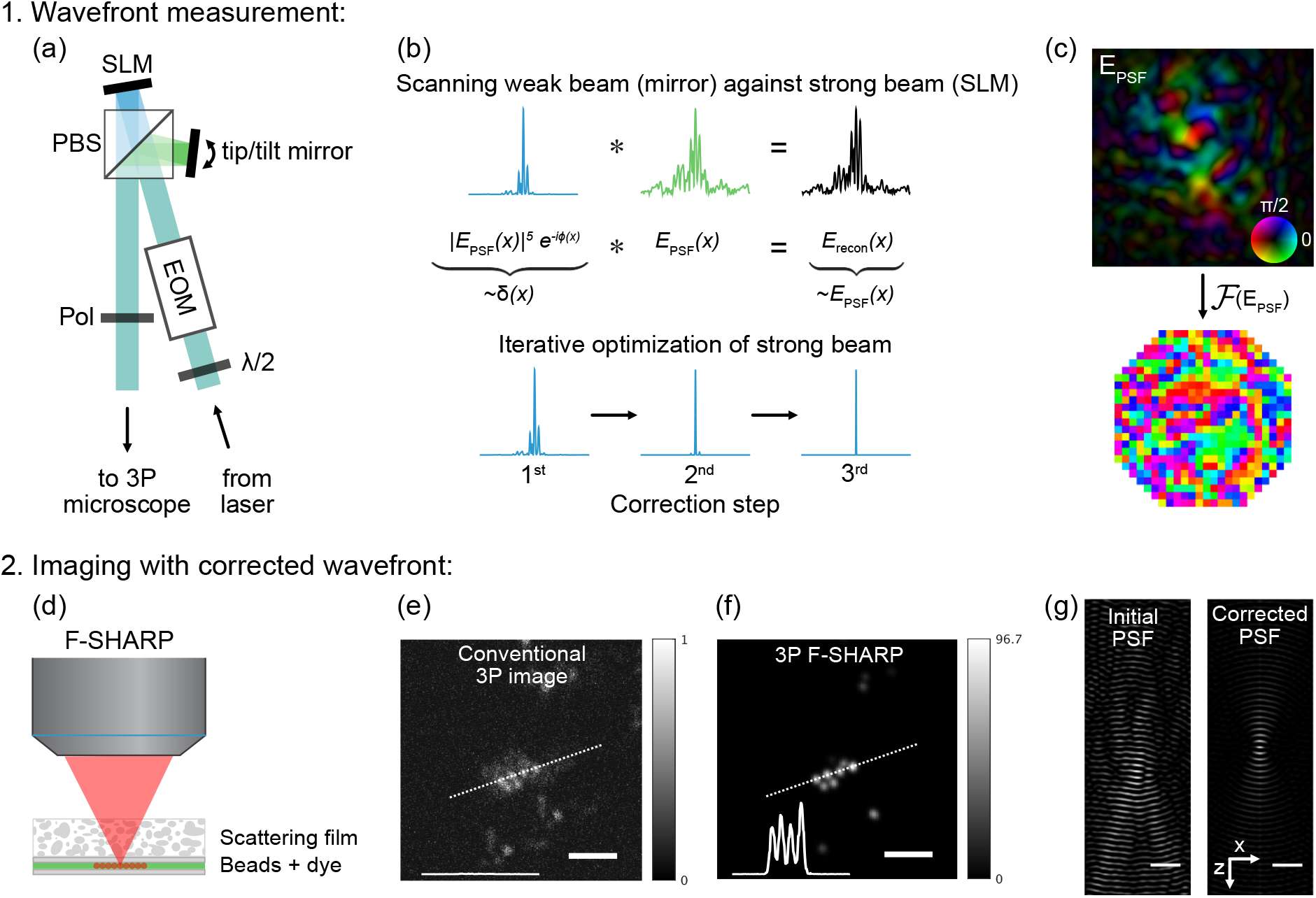
Setup, principle and example measurement of 3P F-SHARP. (a) Schematic of the F-SHARP module, which can be added to a conventional 3P microscope (*λ*/2: half-wave plate, EOM: electro-optical modulator, SLM: spatial light modulator, PBS: polarizing beam splitter, Pol: linear polarizer). (b) Principle of the wavefront measurement: the tip/tilt mirror scans a weak beam against a parked strong beam, which leads to a reconstruction of the E-field PSF at the focal plane *E*_recon_. With each iterative optimization step, the strong beam modulated by the SLM becomes more point-like and the reconstructed field *E*_recon_ becomes closer to the actual *E*_PSF_. (c) From the measured *E*_PSF_ (amplitude encoded in brightness, phase in colormap), the SLM correction pattern (bottom) can be calculated via a Fourier transform. (d) Schematic of the sample. Red fluorescent beads are dispersed in a thin layer of fluorescein. The F-SHARP algorithm only has access to the homogeneous fluorescein signal, while the beads are used for visualization of the signal improvement. (e) 3P images with system correction and (f) with full F-SHARP correction. The intensity profiles along the dashed line are shown as an inset. The images are scaled by the difference of the maxima for better visibility, while the insets are shown to scale. (g) Initial and corrected E-field PSF (real component) plotted along the y-z plane. Both PSFs are scaled to their maximum values to emphasize the differences in spatial confinement. Scale bars 5 µm.

Assuming both beams travel through the same medium and undergo the same scattering (valid within the isoplanatic patch defined by the memory effect (37)), both are scaled versions of *E*_PSF_. With a uniform fluorescent sample and 3P excitation, the resulting image can be described as

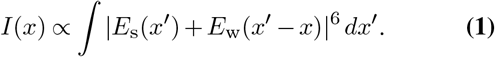

We assume that the intensity of the weak, scanning beam *E*_s_ is much lower than the strong beam and therefore discard all powers of *E*_w_ equal to or larger than two in the algebraic expansion of Eq. 1, as their contribution to the signal can be neglected. This leaves

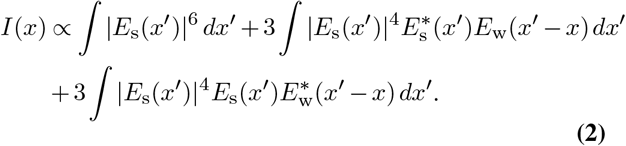

The first term is a uniform background. It can be shown that the second and third terms provide an estimated reconstruction of the scattered E-PSF, *E*_recon_ ≈ *E*_PSF_ and its complex conjugate

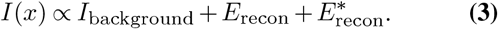

*E*_recon_ can then be isolated from the other terms by a phasestepping scheme (38) using three phase steps.

Assuming *E*_recon_ is an approximation to *E*_PSF_, we can calculate a pattern to correct for the measured aberrations (e.g. via Fourier transform, when the SLM is in the conjugate Fourier plane of the microscope). This correction is then applied onto the SLM, making the strong beam more pointlike. In turn, the estimation of *E*_PSF_ improves. This way, the strong beam iteratively approaches its ideal, diffractionlimited shape. This beam can then be used to image the sample, while the weak beam is blocked by a shutter.

It can be shown that for multiphoton absorption of order *p*, the corrected beam is taken to its (2*p* − 1)th power after every iteration *n* (Supplement 1),

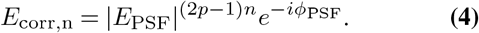

For 3P microscopy (*p* = 3), this predicts convergence to the fifth power of the scattered *E*_PSF_. This is an advantage compared to 2P F-SHARP (*p* = 2), which converges to the third power of the initial *E*_PSF_ after every iteration (34). The full derivation and further details can be found in Supplement 1. The F-SHARP process can be performed with a compact modular addition to any conventional multi-photon microscope (Fig. 1a). The laser (Opera-F, pumped by Monaco, Coherent) provides excitation light at 1300 nm wavelength and 1 MHz repetition rate. The incoming beam is first split and then recombined using a polarizing beamsplitter (with “strong beam” and “weak beam” corresponding to the two polarisation components). A half-wave plate (AHWP05M-1600, Thorlabs) rotates the polarization axis and determines the power ratio between the two beams in this interferometric setup. The “strong beam” propagates towards a segmented deformable mirror (492-SLM, Boston Micromachines Corporation), which is conjugated to the back focal plane of the microscope objective. The “weak beam” is reflected off a tip/tilt mirror (S-331.2SL, Physik Instrumente), which is used to scan this beam in two dimensions. The electro-optic modulator (EOM, EO-AM-NR-C3, Thorlabs) introduces a relative phase shift between the horizontally and the vertically polarised beam (for details on the mechanism, see (35)). Finally, a linear polarizer oriented at 45° allows the two recombined beams to interfere. A detailed setup schematic, as well as a list of components, can be found in Supplement 2.

## Results

### Proof-of-principle

To test the performance of 3P F-SHARP, we covered a thin layer of a mixture of fluorescein (fluorescein sodium salt, Sigma-Aldrich) solution and red-emitting fluorescent beads (1 µm latex beads, amine-modified polystyrene, fluorescent red, Sigma-Aldrich) with a diffuse scattering film (Fig. 1d). Using the feedback signal from this homogeneous dye layer, we determined the scattered *E*_PSF_ within two 3P F-SHARP iterations. The laser power was gradually lowered after each iteration to maintain a comparable signal intensity. To measure the improvement, we then imaged the fluorescent beads with conventional 3P microscopy (corrected for system aberrations) (Fig. 1e). We then imaged the same field of view using identical laser power (0.9 mW) with the full 3P F-SHARP correction (Fig. 1f). In the conventional 3P image, the beads are barely visible above the noise level. As the 3P excitation probability scales with the third power of the excitation power density, out-of-focus excitation is low and the beads are still recognizable even with a strongly aberrated PSF. However, the signal intensity is increased by nearly two orders of magnitude using the full 3P F-SHARP correction. The corresponding E_PSF_ and the SLM correction pattern are shown in Fig. 1c. By propagating this electric field along the z-axis using the angular spectrum method (39), we obtain the 3D E-field PSF before and after correction (Fig. 1g). To calculate the corrected PSF, we assume that the phase can be corrected perfectly by the SLM, and only amplitude variations remain. The corrected PSF along the y-z plane shows that a nearly diffraction limited PSF can be recovered.

### Convergence on a bulk fluorescent sample

Most wavefront shaping techniques rely on feedback of a guidestar or nonlinear feedback from a confined region (24). For example, Katz et al. (40) optimize the total 2P-excited signal for wavefront shaping through highly scattering layers. They note that this method works on planar fluorescent samples but does not converge on a homogeneous three-dimensional sample. Intuitively, this can be understood through the coupling of lateral and axial resolution: a more point-like focus increases the maximum signal, but at the same time shrinks the excitation volume axially, keeping the total fluorescence constant (41). The total 2P excited fluorescence therefore ceases to be a good measure of focus quality in thick fluorescent samples. For 3P excitation, however, the increased peak power of a smaller focus outweighs the effects of the shrinking excitation volume (40, 42). Therefore, we expect 3P F-SHARP to converge even in a homogeneous threedimensional fluorescent object.

To illustrate this, we can consider a Gaussian beam focused to a waist *w*_0_, whose focus volume can be approximated by a cylinder with length 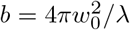 In this case, the total generated signal is proportional to (40)

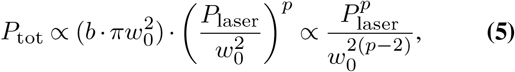

with the total excitation power *P*_laser_. For *p* = 2, this is independent of beam waist diameter. Only for *p >* 2, the total signal increases with a tighter focus. An exact solution comparing 2P and 3P signal per pulse can be found in Wang et al. (42).

To demonstrate the different convergence in bulk fluorescence, we compared the performance of 2P and 3P F-SHARP on a thin dye layer (Fig. 2a,b) and in a dye volume (Fig. 2c,d), both covered by a strongly scattering film. For a fair comparison between 2P and 3P F-SHARP, we aimed to have the same scattered E_PSF_ and therefore use the same excitation wavelength in both cases. Thus, we used a mixture of two dyes which can both be excited at 1300 nm, but with either 2P or 3P excitation: fluorescein (Sigma-Aldrich) for 3P excitation at 1300 nm and Alexa Fluor 680 (Invitrogen) for 2P excitation at 1300 nm (43).

**Fig. 2.**
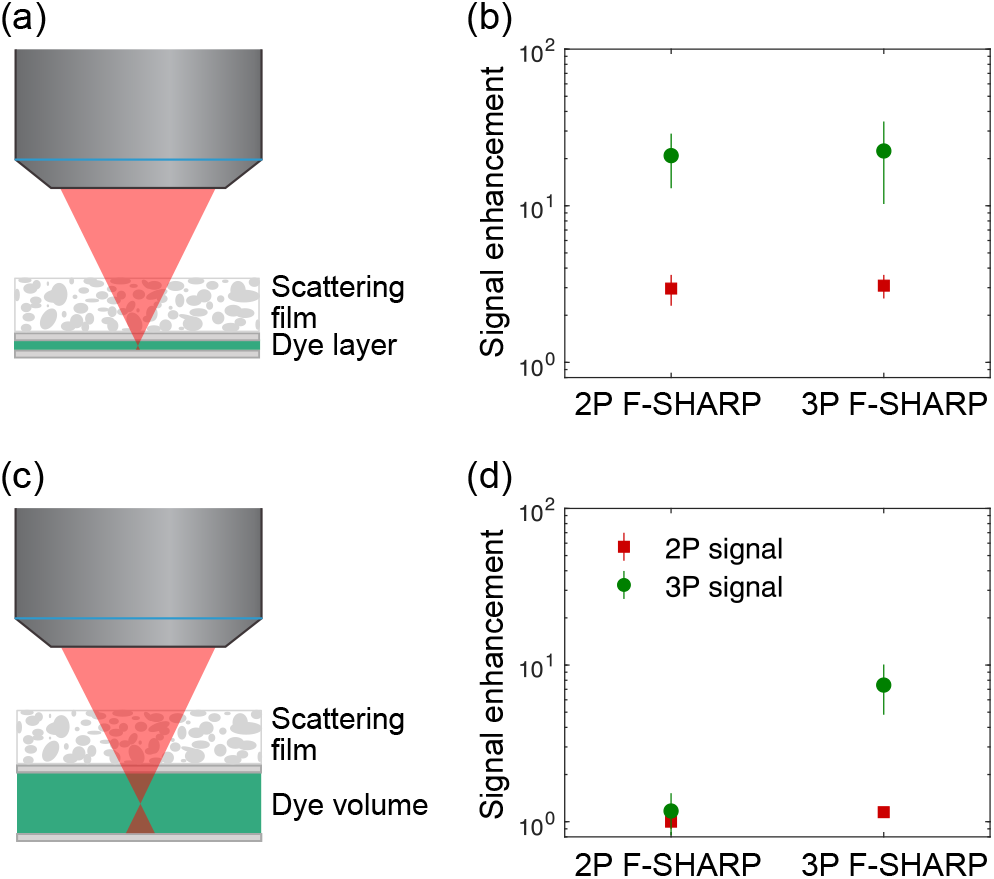
3P F-SHARP does not depend on the distribution of fluorescence. (a) Schematic of the sample geometry to compare the performance on an approximately two-dimensional sample. (b) Comparison of signal improvement using 2P or 3P F-SHARP. Shown are mean and standard deviation of 8 values each. Both the 2P signal improvement (red) and the 3P signal improvement (green) are comparable for 2P and 3P F-SHARP. Due to the higher order nonlinearity, the 3P signal improvement is larger. (c) Sample geometry to evaluate the performance in an extended sample. (d) Comparison of signal improvement for both F-SHARP methods in a 3D sample. Only 3P F-SHARP converges and enhances the signal.

We investigated whether the signal intensity within the isoplanatic patch increases in response to either 2P F-SHARP or 3P F-SHARP corrections. For each correction pattern, we measured both 2P (red channel) and 3P signal (green channel) improvement. For the same change in E_PSF_, the 3P signal improves by a larger factor due to the higher order nonlinear excitation (11), as increased peak power nonlinearly enhances the total signal. Positions (*n* = 8 per category) were selected randomly. On each position, one 2P F-SHARP and one 3P F-SHARP measurement were performed, with alternating order to avoid any systematic effects due to photo-bleaching. Excitation power was chosen such that the signal was comparable or favorable for 2P F-SHARP (3 mW for 3P excitation of fluorescein, <1 mW for 2P excitation of Alexa Fluor 680). Fig. 2b shows that on a thin dye layer with thickness on the order of the axial resolution, both 2P and 3P FSHARP perform well.

In contrast, 2P F-SHARP fails to converge on a homogeneous three-dimensional dye volume, and both 2P and 3P signal stay almost constant (Fig. 2d). In contrast, 3P F-SHARP converges even on a thick homogeneous sample with strong scattering. As expected, the 3P F-SHARP correction leads to an increased 3P signal, but the 2P signal also does not increase even for a correction estimated by 3P F-SHARP, as a tighter focus does not increase the total fluorescence.

### Fast convergence to fifth power of *E*_PSF_

As shown above, the scattered *E*_PSF_ should be taken to its fifth power during each step of the iterative 3P F-SHARP process (for details, see Supplement 1). This is faster than for 2P F-SHARP, where a convergence to the third power is expected. To show this experimentally, we used the same setup as for the proof-of-principle experiments (Fig. 3a). The F-SHARP correction was performed on a thin layer of fluorescein solution, covered by a scattering film. A sparse distribution of red fluorescent beads was then used to visualize the 3P PSF at each iteration (Fig. 3b). Each FOV only contained a single bead.

**Fig. 3.**
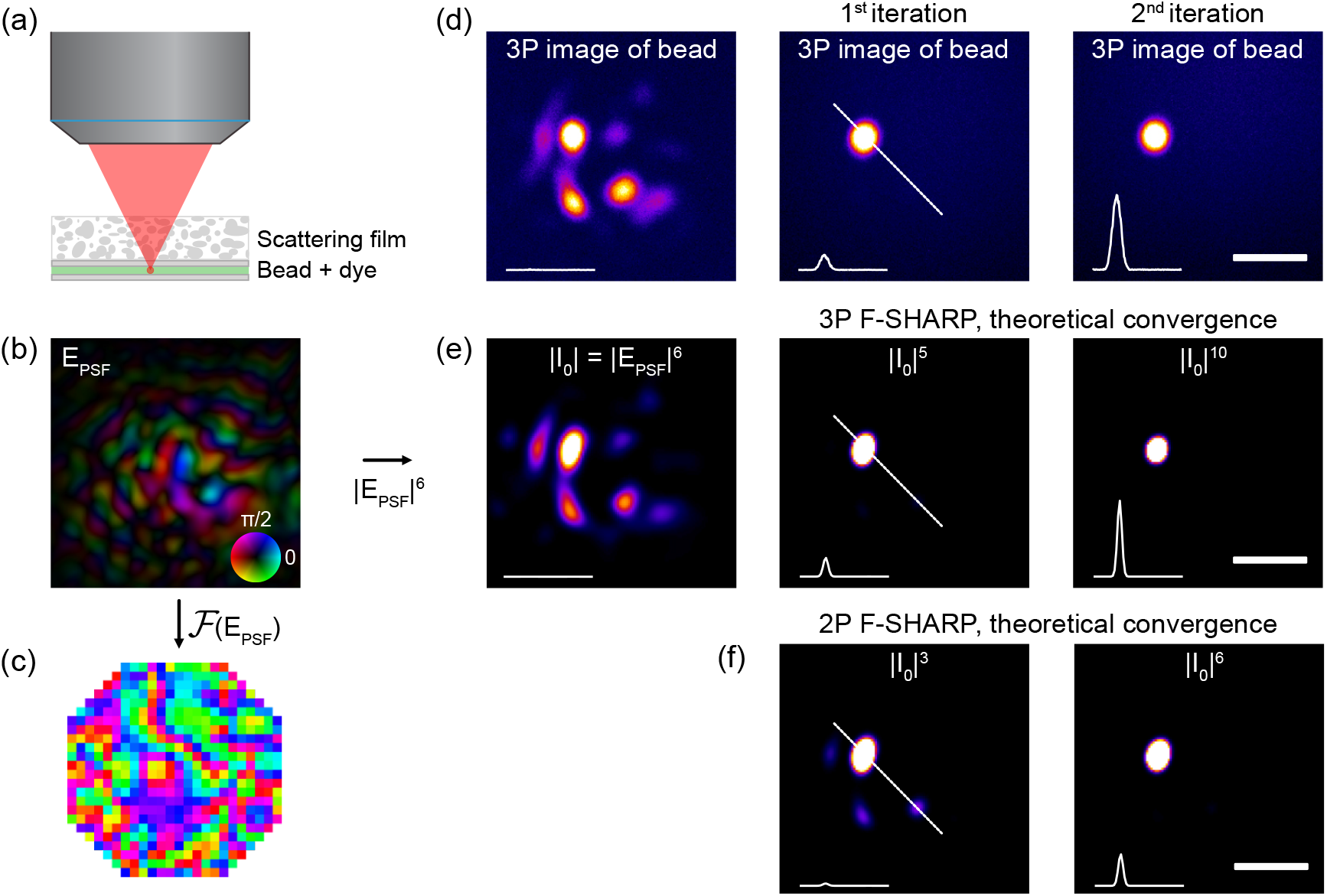
3P F-SHARP converges to the fifth power of the scattered *E*_PSF_. (a) F-SHARP measurements are performed on a thin layer of fluorescein covered by a scattering film. The corresponding 3P PSF is observed using a single, red fluorescent bead. (b) Scattered *E*_PSF_ estimated after two iterations of F-SHARP and (c) corresponding correction pattern. (d) Initial 3P image of bead without correction, showing strong aberrations. Within two correction steps, the 3P PSF becomes point-like and the signal strongly increases. The insets show profiles along the dashed line. Images are presented saturated to visualize weaker side lobes, while insets are shown to scale. (e) By taking the sixth power of the estimated scattered *E*_PSF_, the 3P point spread function *I*_0_ can be calculated. This corresponds well to the directly measured PSF in (d). The following images show the theoretically predicted convergence to the fifth power of *|I*_0_*|*. (f) For 2P F-SHARP, convergence to the third power is expected. The measured PSFs in (b) show that 3P F-SHARP converges faster. Scale bars 3 µm.

Over the course of two iterations of 3P F-SHARP, the PSF converges from a heavily scattered pattern with multiple peaks to a single peak with strongly enhanced intensity (see insets of profiles shown to scale). Fig. 3b shows the estimated *E*_PSF_ after the second iteration. From this, we can calculate the SLM correction pattern (Fig. 3c) and the initial 3P PSF *I*_0_, with *I*_0_ = |*I*_PSF_ |^3^ = |*E*_PSF_| ^6^ (Fig. 3e). This estimation is very similar to the measured shape in Fig. 3d, which indicates that the F-SHARP measurement precisely estimates the scattered *E*_PSF_ after only two iterations.

Furthermore, we calculate the theoretically predicted convergence behaviors for 3P F-SHARP (Fig. 3e) and 2P F-SHARP (Fig. 3f). The measurements are in good agreement with the 3P F-SHARP prediction. In contrast, for the theoretically expected convergence of 2P F-SHARP, side lobes of the PSF are still clearly visible after the first iteration. This faster convergence of 3P F-SHARP reduces the photon budget and time requirements for wavefront measurement.

### Application to *in vivo* imaging

To demonstrate potential applications of 3P F-SHARP, we use it to enhance the signal acquired from GFP-labeled pyramidal neurons in an anesthetized mouse *in vivo* (Fig. 4a, see Supplement 3 for details on the preparation). 3P F-SHARP was performed with the strong beam parked on the soma, converging after two iterations. The corresponding SLM correction pattern and predicted three-dimensional PSFs before and after correction are shown in Fig. 4c and e. We compare maximum intensity projections (MIPs) of 11 planes at 1 µm distance, five below and five above the focal plane of the correction (Fig. 4b). Each plane consists of an average of five frames motion-corrected with NorRM-Corre (44). The corrected image exhibits an approximately eight-fold increase in signal (Fig. 4d), as well as improved resolution. After 3P F-SHARP correction, dendritic spines can be clearly distinguished.

**Fig. 4.**
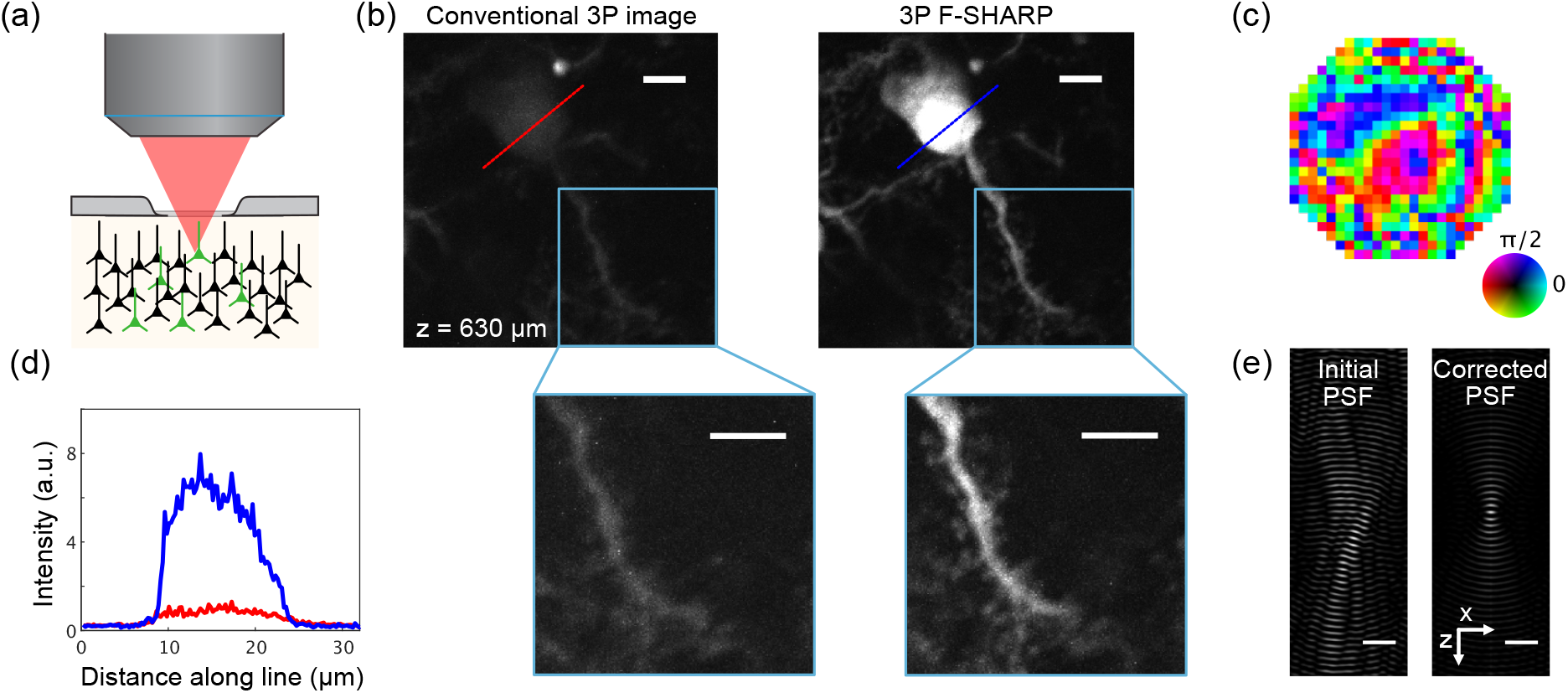
Scattering compensation inside a living mouse brain. (a) Schematic of *in vivo* mouse brain imaging. Imaging is performed in an anesthesized mouse through a craniotomy. (b) Comparison of conventional 3P image (only system correction) and 3P F-SHARP corrected image (maximum intensity projection of 11 planes, *±* 5 µm) of a GFP-expressing pyramidal neuron 630 µm below the brain surface. Insets show magnified views of dendrite and spines. Brightness is shown on the same scale, with the corrected image partially saturated. Scale bars 10 µm. Excitation power at the surface was 16 mW at 1 MHz repetition rate. (c) Correction pattern. (d) Profiles along dashed lines in (b) show signal increase along the soma. Traces were normalized to the maximum of the uncorrected trace, noise-filtered with a 5-pixel moving median filter. (e) Three-dimensional electric field PSFs (real component) plotted along the y-z plane, both scaled to their maxima. Scale bars 5 µm.

## Discussion

In this work, we presented 3P F-SHARP, a scattering compensation method based on 3P microscopy. In addition to the improved penetration depth of long wavelength 3P excitation, the higher order nonlinearity leads to qualitatively different performance. In contrast to 2P wavefront shaping techniques, 3P F-SHARP does not need to rely on the distribution of fluorescence: While 2P F-SHARP fails to improve the point spread function with feedback from a three-dimensional homogeneous dye sample, 3P F-SHARP converges and improves the signal. This means that 3P F-SHARP can give an estimate of the scattered *E*_PSF_ even in the absence of any contrast, which can be beneficial in densely labeled regions.

Additionally, we mathematically predicted that 3P F-SHARP converges faster than comparable techniques with 2P excitation and confirmed this experimentally. Fast convergence is essential for *in vivo* experiments, as it accelerates the wavefront optimization process and requires fewer photons.

In theory, phase stepping requires only two measurements. Usually, more measurements are taken to get a more robust estimate. Here, we show that reducing the phase steps from four to three is possible and still works reliably.

Each iteration in the presented experiments takes approximately three seconds. However, this imaging speed could be increased further. The scanning speed was limited to 200 Hz line rate by the two-axis mirror. With 1 kHz line rate achievable with a higher power amplifier and the minimum number of measured modes, each iteration would only take 72 ms. Usually, two to three iterations were sufficient even for highly scattered PSFs (compare Fig. 1 and Fig. 3).

A further increase of the imaging depth would be possible, as the system is mainly limited by available laser power. Reduced transmission of optical components for 3P compared to 2P excitation wavelengths, as well as the linear polarizer necessary for interference of weak and strong beam, lead to a low power efficiency of the system. The F-SHARP module itself has a power transmission efficiency of approximately 35 %, dominated by the 50 % loss at the linear polarizer. The maximum power available under the objective is around 25 mW. However, higher power laser systems have become available and could be used with 3P F-SHARP. Custom coatings for optical components could improve the overall transmission. Additionally, both power transmission and the number of corrected modes could potentially be improved by limiting the incident NA of the microscopy system, as suggested by Jin et al. (45).

In summary, 3P F-SHARP is a scattering compensation method with fast convergence on samples with arbitrary fluorescence structure. With sufficient excitation power, we expect that it will push the depth limits of optical imaging.

## ACKNOWLEDGEMENTS

The authors thank M. Hoffmann and M. Kadobianskyi for helpful discussions and for critically reviewing the manuscript. The authors also thank A. Ender, A. Groneberg, J. Henninger and D. Owald for help with preliminary experiments. This preprint is formatted using the Henriques lab bioRxiv template.

