## Supplemental document for "Deep tissue scattering compensation with three-photon F-SHARP"

### 1. MATHEMATICAL BASIS OF 3P F-SHARP

Here, we derive the mathematical basis for 3P F-SHARP. Assuming a uniform fluorescent sample, we can describe the measured intensity signal as a superposition of two beams: the "strong beam" $E_s$, parked in the center of the field of view and a scanning "weak beam" $E_w$. The measured 3P intensity as a function of scanning position $x$ can then be described as

$$I(x) \propto \int |E_s(x') + E_w(x'-x)|^6 \, dx'. \tag{S1}$$

At the beginning of the F-SHARP measurement, we assume that both beams experience the same scattering. This means that both $E_s$ and $E_w$ are scaled versions of the scattered $E_{PSF}$. This assumption is valid within the memory effect range (the isoplanatic patch), where $E_w$ does not change its shape depending on the scanning position and thus only depends on the difference $x' - x$.

We can then expand Eq. S1 with $a = E_s$ and $b = E_w$,

$$\begin{aligned} |a+b|^6 =& |a|^6 + |b|^6 + |a|^4(3ab^* + 3a^*b + 9|b|^2) + |b|^4(9|a|^2 + 3ab^* + 3a^*b) + ... \\ &... + a^{*3}b^3 + a^3b^{*3} + 3(|a|^2 + |b|^2)(a^2b^{*2} + a^{*2}b^2) + 9|a|^2|b|^2(a^*b + ab^*). \end{aligned} \tag{S2}$$

If |a| ≫ |b| ($|E_s| \gg |E_w|$), we can neglect all terms that contain b to power two or higher, which leaves

$$|a+b|^6 \approx |a|^6 + 3|a|^4(ab^* + a^*b). \tag{S3}$$

The measured intensity can thus be approximated as

$$\begin{aligned} I(x) \propto & \int |E_s(x')|^6 \, dx' \\ & + 3\int |E_s(x')|^4 E_s^*(x') E_w(x'-x) \, dx' \\ & + 3\int |E_s(x')|^4 E_s(x') E_w^*(x'-x) \, dx'. \end{aligned} \tag{S4}$$

The first term can be described as a uniform background. In the following, we will show that the second and third terms provide an estimate for the scattered E-field PSF, $E_r \approx E_{PSF}$ and its complex conjugate:

$$I(x) \propto I_{background} + E_r + E_r^*. \tag{S5}$$

We start by assuming a plane wave at the back focal plane of the objective, which then travels both through a lens (the objective) and an inhomogeneous medium. The resulting field at the focal plane is the scattered $E_{PSF}$. Equivalently, one could consider a modulated (non-planar) input wavefront at the back focal plane, which only travels through the lens and not through the medium, but still results in $E_{PSF}$ at the focal plane. This input wavefront $E_{in}$ is then related to $E_{PSF}$ via a Fourier transform

$$E_{in} = \mathcal{F}(E_{PSF}). \tag{S6}$$

We can now use the pattern $P_{SLM}$ applied to the SLM, which is in a conjugate plane to the back aperture, to correct for these aberrations. The field at the back aperture can then be expressed as

$$E_{BA} = P_{SLM} \cdot \mathcal{F}(E_{PSF}). \tag{S7}$$

If we had exact knowledge of $E_{PSF}$, we could apply

$$P_{SLM} = \mathcal{F}(E_{PSF})^* \tag{S8}$$

and due to the time reversal property of light in the absence of absorption, all aberrations would be cancelled,

$$\mathcal{F}(E_{\mathrm{PSF}})^* \cdot \mathcal{F}(E_{\mathrm{PSF}}) = 1. \tag{S9}$$

Instead, we will show that we can use $E_{\mathrm{r}}$ as an approximation to $E_{\mathrm{PSF}}$,

$$E_{\mathrm{r}}(x) \propto \int |E_{\mathrm{s}}(x')|^4 E_{\mathrm{s}}^*(x') E_{\mathrm{w}}(x'-x) dx'. \tag{S10}$$

To do so, we first substitute $x \to -x$, which corresponds to flipping the field

$$E_{\mathrm{r}}(-x) \propto \int |E_{\mathrm{s}}(x')|^4 E_{\mathrm{s}}^*(x') E_{\mathrm{w}}(x'+x) dx'. \tag{S11}$$

This enables us to use the cross-correlation Fourier theorem to calculate

$$\mathcal{F}\{E_{\mathrm{r}}(-x)\} \propto \mathcal{F}\{|E_{\mathrm{s}}(x)|^4 E_{\mathrm{s}}^*(x)\}^* \cdot \mathcal{F}\{E_{\mathrm{w}}(x)\} \tag{S12}$$

and

$$\mathcal{F}\{E_{\mathrm{r}}(-x)\}^* \propto \mathcal{F}\{|E_{\mathrm{s}}(x)|^4 E_{\mathrm{s}}^*(x)\} \cdot \mathcal{F}\{E_{\mathrm{w}}(x)\}^*. \tag{S13}$$

Instead of Eq. S8, we can then apply

$$P_{\mathrm{SLM}} = \mathcal{F}(E_{\mathrm{r}}(-x))^* \tag{S14}$$

on the SLM. At the back aperture, we then get

$$\begin{aligned} E_{\mathrm{BA}} &= P_{\mathrm{SLM}} \cdot \mathcal{F}(E_{\mathrm{PSF}}) \\ &= \mathcal{F}\{|E_{\mathrm{s}}(x)|^4 E_{\mathrm{s}}^*(x)\} \cdot \mathcal{F}\{E_{\mathrm{w}}(x)\}^* \cdot \mathcal{F}(E_{\mathrm{PSF}}). \end{aligned} \tag{S15}$$

As $E_{\mathrm{w}}(x) \approx E_{\mathrm{PSF}}(x)$, we can use Eq. S9 to show that

$$E_{\mathrm{BA}} = \mathcal{F}\{|E_{\mathrm{s}}(x)|^4 E_{\mathrm{s}}^*(x)\}. \tag{S16}$$

The field at the focal plane after the first iteration $n = 1$, $E_{\mathrm{corr,n=1}}(x)$, is then the inverse Fourier transform of $E_{\mathrm{BA}}$

$$E_{\mathrm{corr,n=1}}(x) \approx |E_{\mathrm{s}}(x)|^4 E_{\mathrm{s}}^*(x). \tag{S17}$$

In this first iteration, $E_{\mathrm{s}}(x) \approx E_{\mathrm{PSF}}(x)$ and thus

$$E_{\mathrm{corr,n=1}}(x) \approx |E_{\mathrm{PSF}}(x)|^5 e^{-i\phi_{\mathrm{PSF}}(x)}. \tag{S18}$$

In subsequent iterations, the strong beam has already been corrected by the previous iteration, $E_{\mathrm{s,n}}(x) = E_{\mathrm{corr,n-1}}(x)$, and thus

$$E_{\mathrm{corr,n}}(x) = |E_{\mathrm{PSF}}(x)|^{5n} e^{-i\phi_{\mathrm{PSF}}(x)}. \tag{S19}$$

This shows that the $E_{\mathrm{corr}}(x)$ will converge to the fifth power of the previous E-field PSF in every iteration.

After every step, the correction (Eq. S14) is applied on the SLM, making the strong beam more point-like. In turn, the estimation of $E_{\mathrm{PSF}}$ improves. This way, the strong beam iteratively approaches its ideal, diffraction limited shape. After the algorithm has converged, this beam can be used to image the sample.

Thus, we need to determine $E_{\mathrm{r}}$ to iteratively turn the strong beam into a diffraction-limited focus, with each iteration taking the previous field to its fifth power. To do so, we can isolate $E_{\mathrm{r}}$ from the other terms in Eq. S4 via a phase-stepping scheme. By setting a phase difference $\Delta\varphi_i$ between the two beams, we get

$$\begin{aligned} I_k(x) \propto &\int |E_{\mathrm{s}}(x')|^6 \, dx' \\ &+ 3 \int |E_{\mathrm{s}}(x')|^4 E_{\mathrm{s}}^*(x') E_{\mathrm{w}}(x'-x) e^{-i\Delta\varphi_k} \, dx' \\ &+ 3 \int |E_{\mathrm{s}}(x')|^4 E_{\mathrm{s}}(x') E_{\mathrm{w}}^*(x'-x) e^{i\Delta\varphi_k} \, dx'. \end{aligned} \tag{S20}$$

The standard solution for a four-step phase-stepping scheme, which was also used for 2P F-SHARP, is

$$E_r = (I_0 - I_\pi) + i(I_{\pi/2} - I_{3\pi/2}), \tag{S21}$$

where the phase difference between the two beams is $\Delta\varphi_k = 0, \pi/2, \pi, 2\pi/3$. However, in this implementation of 3P F-SHARP, the phase range is limited to $\pm\pi/2$ by the EOM used to introduce the phase shift. Additionally, fewer phase steps are desirable to determine the correction faster.

We start with Eq. S20 in a simplified form

$$I_k(x) = c + d(x)e^{-i\Delta\varphi_k} + d^*(x)e^{i\Delta\varphi_k}, \tag{S22}$$

where

$$c = \int |E_\mathrm{s}(x')|^6 \, dx' \tag{S23}$$

and

$$d(x) = 3\int |E_\mathrm{s}(x')|^4 E_\mathrm{s}^*(x') E_\mathrm{w}(x' - x)\, dx'. \tag{S24}$$

The aim is to isolate $d(x)$ with $m$ uniformly spaced phase-shifts $\Delta\varphi_k$. We can find a general solution for $m$ phase steps by solving the set of equations

$$\sum_{k=0}^{m-1} z_k I_k = \lambda d, \tag{S25}$$

with $\lambda \neq 0$.

In this implementation, we chose $m = 3$ and $\Delta\varphi_k = -\pi/2, 0, \pi/2$, limited by the range of the electro-optic modulator (EOM). To center the phase shifts around zero, necessary for the EOM, we introduce a global phase shift of $e^{i\frac{5}{4}\pi}$, which does not influence the F-SHARP measurement. This leads to the solution $z_{-\pi/2} = i$, $z_0 = -(i+1)$ and $z_{\pi/2} = 1$, with $\lambda = -2$.

This means that we can determine $E_r$ from the intensities measured during the three phase steps via

$$E_r \propto i \cdot I_{-\pi/2} - (i+1)I_0 + I_{\pi/2}. \tag{S26}$$

### 2. EXPERIMENTAL SETUP

The F-SHARP setup consists of a laser unit with dispersion compensation and power control, an F-SHARP module and a regular, home-built multiphoton microscope (Fig. S1). Components are listed below (Table S1).

In short, the laser output first passes through a dispersion compensation unit based on a single prism. The power is adjusted using a motorized half-wave plate and a polarizing beam splitter. A second half-wave plate sets the power ratio between horizontally and vertically polarized light, and thus between strong beam and weak beam. A telescope (L1/L2) decreases the beam diameter for it to pass the electro-optic modulator (EOM) without clipping. A second telescope (L3/L4) expands the beam to fill the aperture of the spatial light modulator (SLM). A polarizing beam splitter separates and recombines horizontally and vertically polarized light. The strong beam is reflected towards the SLM, while the weak beam is transmitted towards a tip/tilt mirror. A linear polarizer (LP) allows the two beams to interfere. A third telescope (L5/L6) adjusts the beam diameter to fill the galvo mirrors. It also conjugates the SLM plane to the first galvo, and thus to the back aperture of the objective. Three scan lenses, two galvo mirrors and a tube lens form a conventional scanning microscope. Fluorescence is detected in two channels with photomultiplier tubes (PMTs).

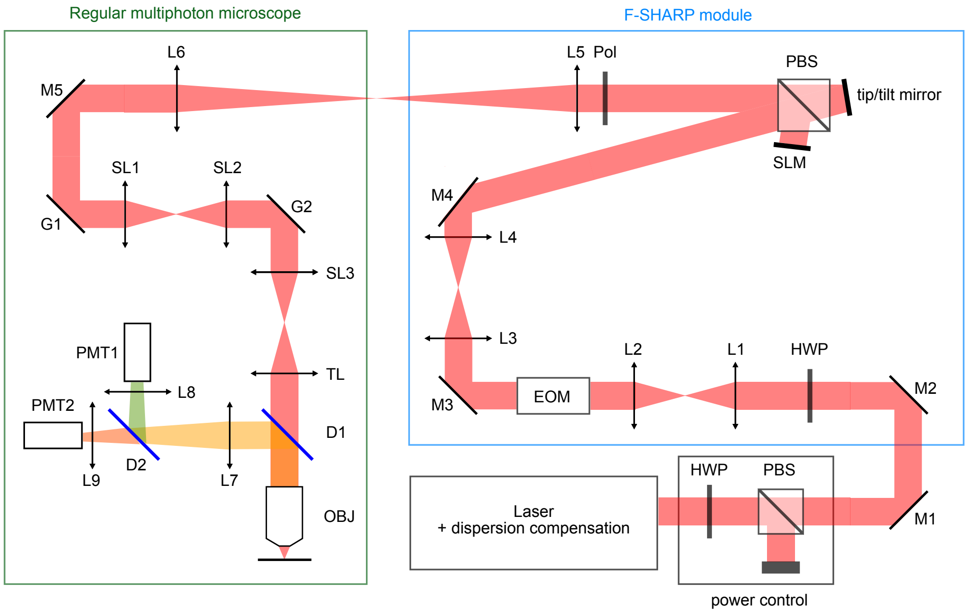


**Fig. S1. Detailed experimental setup.** The F-SHARP module (blue) can be used as a modular add-on to a regular multiphoton microscope (green). HWP: half-wave plate, PBS: polarizing beam splitter, M: mirror, L: lens, EOM: electro-optic modulator, SLM: spatial light modulator, Pol: linear polarizer, G: galvo mirror, SL: scan lens, TL: tube lens, D1: primary dichroic, OBJ: objective, D2: secondary dichroic, PMT: photomultiplier tube. A full list of optical components can be found in Table 1.

| | |
|---|---|
| Laser | Opera-F, Light Conversion/Coherent pumped by Monaco, Coherent |
| HWP | AHWP05M-1600, Thorlabs |
| PBS | PBS124, Thorlabs |
| M1-M5 | PF10-03-P01, Thorlabs |
| L1 | AC254-250-C-ML, Thorlabs |
| L2 | AC127-075-C-ML, Thorlabs |
| EOM | EO-AM-NR-C3, Thorlabs |
| EOM amplifier | A800, FLC Electronics |
| L3 | AC127-025-C-ML, Thorlabs |
| L4 | AC254-150-C-ML, Thorlabs |
| SLM | 492-DM, Boston Micromachines Corporation |
| Tip/tilt mirror | S-331.2SL, Physik Instrumente |
| Pol | LPNIRC050-MP2, Thorlabs |
| L5 | AC254-200-C-ML, Thorlabs |
| L6 | AC254-125-C, Thorlabs |
| SL1/2 | LSM05, Thorlabs |
| SL3 | SL50-3P, Thorlabs |
| TL | AC508-200-C-ML |
| D1 | DMLP650L or DMLP805L, Thorlabs |
| OBJ | Nikon LWD 25x/1.10W |
| L7 | LA1050-A, Thorlabs |
| D2 | F39-562, AHF Analysentechnik |
| L8/9 | ACL2520U-A, Thorlabs |
| PMT1/2 | H10770PA-40 MOD, Hamamatsu |
| Filter 1 (PMT1) | BrightLine® Fluorescence Filter 525/50, Semrock |
| Filter 2 (PMT2) | BrightLine® Fluorescence Filter 607/70, Semrock or FB700-40, Thorlabs |

**Table S1.** List of components

### 3. MOUSE PREPARATION

All experiments were performed according to protocols approved by the Berlin Animal Ethics committee (Landesamt für Gesundheit und Soziales, LAGeSo) and complied with the European animal welfare law. 5 week old C57BL/6J wild-type mice were used in this study. Mice were anaesthetized with ketamine (100 mg $kg^{-1}$)/xylazine (10 mg $kg^{-1}$) and kept on a thermal blanket to maintain body temperature. Ophthalmic ointment was used to protect the animal's eyes during the surgery. The scalp and periosteum were carefully removed and a 4-5 mm craniotomy was made over the right S1 centered at stereotaxic coordinates, 1.5 mm posterior and 2.2 mm lateral from bregma. For GFP expression, AAV2/9–CAG–GFP (Charité Vector Core Facility) was injected through a glass pipette (tip diameter, 10-30 µm). A volume of 50 nL of virus was injected at 50 nL/min at three depths at 900, 700 and 500 µm deep from the pial surface. After injection, the craniotomy was sealed with a 4 mm glass coverslip with cyanoacrylate glue and dental cement. Finally, a light-weight head-post was fixed on the skull in the left hemisphere with light-curing adhesives and dental cement. Head-fixed imaging experiments began 3 weeks after the virus injection under ketamine/xylazine anesthesia.
